# The interaction between a parasite and sub-optimal temperatures contributes to honey bee decline

**DOI:** 10.1101/2021.07.22.453350

**Authors:** Davide Frizzera, Laura Andreuzza, Giulia Boaro, Mauro D’Agaro, Simone Del Fabbro, Virginia Zanni, Desiderato Annoscia, Francesco Nazzi

## Abstract

Global insect decline and, in particular, honey bee colony losses are related to multiple stress factors, including landscape deterioration, pollution, parasites and climate change. However, the implications of the interaction among different stress factors for insect health are still poorly understood; in particular, little is known on how challenging environmental conditions can influence the impact of parasites. Here we exploited the honey bee as a model system to approach this problem and carried out extensive lab and field work aiming at assessing how suboptimal temperatures and parasitic challenges can alter the homeostatic balance of individual bees and the whole colony, leading to individual death and colony collapse.

We found that mite infestation further than increasing the mortality of bees, induces an anorexia that in turn reduces the capacity of bees to thermoregulate, thus exposing them to the detrimental effect of lower temperatures. This, in turn, has important implications for the colony as a whole. The results highlight the role that abiotic factors can have in shaping the effect of parasitic challenges on honey bees. Furthermore, the multilevel and holistic approach adopted here can represent a useful template for similar studies on other insect species, which are particularly urgent in view of climate change and the continuous pressure of natural and exotic parasites on insect populations.

## 1. Introduction

In recent years, large losses of honey bee colonies have become a global issue ^1,2,3^, causing justified concern in view of the essential pollination service provided by this species^4^. At the same time, other reports revealed similar problems affecting wild bees^5^, confirming previous studies dealing with other insects’ taxa^6^. In 2017, a long population monitoring study carried out in Germany highlighted a 76% decline in flying insect biomass^7^; more recently, another long-term study in the rain forests of Puerto Rico reported biomass losses above 78% for ground-foraging and canopy-dwelling arthropods^8^. Overall, these latter and other studies indicate that several insect taxa are experiencing severe losses^6^, which, in view of the ecological role played by insect in most terrestrial ecosystems, require urgent and careful consideration.

It has been suggested that such declines are caused by a number of potential stressors including habitat loss, pollution, parasites and climate change^6^. However, despite none of these factor comes in isolation, our knowledge about the possible combined effect of more stressors on insect populations is dramatically scarce^9^; in particular, the combination between biotic and abiotic stressors, like parasites and adverse environmental conditions, respectively is still largely unexplored, although this latter case appears particularly important in view of the growing importance of climate change in shaping the already complex interactions within the ecosystems. This lack of knowledge is not surprising in view of the complexity of the necessary multifactorial studies; for example, a recent literature survey of insect studies in which different classes of stressors were manipulated in a full-factorial manner, produced only 133 studies covering 24 stressor pairs^9^. Under this respect, honey bees offer a unique opportunity in view of the detailed knowledge of their biology, the large suite of molecular tools available since the sequencing of the honey bee genome^10^, and the relative ease of access and manipulation^11^, notwithstanding the opportunity of investigating effects at both the individual and colony level.

In the northern hemisphere, where honey bee colony losses are reported, most of them occur during the autumn-winter period ^12,13,14^. In fact, monitoring programs in the US highlighted a higher mortality of bee colonies in northern states^15^, and, in some cases, a correlation was found between winter temperature and colony losses^16^. To our knowledge, in Europe, where extensive surveys of colony losses were carried out, a similar pattern has not been reported so far, although published data support the hypothesis that colony losses are somewhat higher in northern Europe as compared to southern European countries^14^.

It has been shown that colony losses are related to the progressive build-up of viral infections promoted by the increasing *Varroa destructor* infestation^17^. In fact, both mite infestation and deformed wing virus (DWV) prevalence and abundance gradually increase along the season, peaking at the end of Summer, when thousands of mites can be present in each colony, DWV prevalence reaches 100% and viral load in bees can be as high as 10^15^-10^18^ viral particles per bee^17^. This in turn, causes increased honey bee mortality, leading to the progressive weakening of the colony, which eventually collapse during autumn or the following winter^17^. Concurrently, under temperate climatic conditions, a more or less marked decrease of temperature, according to latitude and continentality, is also observed^18^. However, regardless of the external fluctuations, nest temperature is constantly maintained around 34-35 °C^19,20,21^; this, under lower external temperatures, is made possible by the capacity of a conveniently large cohort of bees to warm up their thorax after consuming an adequate supply of honey^22,23^. Since both the number of bees involved in thermoregulation and their efficiency, as well as the supply of honey could be influenced by mite infestation, we hypothesized that the capacity of bees to maintain a convenient nest temperature can be impaired as a result of the increasing mite infestation and asked if and how this can influence the conditions of the bee colony, further aggravating the already negative impact of the parasite. To this aim, we studied the effect of an increasing mite infestation and a concurrent low temperature on individual bees and bee colonies. In general, we aimed at understanding how the influence of a biotic challenge, like a parasitic mite, can be shaped by an abiotic factor, like environmental temperature, both at the individual and colony level. This way we wanted to significantly enlarge the growing body of research about the effects of interacting stress factors on honey bee health^24,25^, eventually including an environmental factor that has been overlooked so far. In doing so we tried to provide a useful reference for similar studies with other insect species.

## 2. Results

### 2.1. Colony conditions according to mite infestation

To study the effect of the decreasing environmental temperature on honey bee colonies exposed or not to the parasitic mite *V. destructor*, we established two groups of hives, one of which was treated with acaricides throughout the trial, to maintain mite infestation at the lowest possible level, while the other was left untreated until October.

As expected, a high number of mites, largely exceeding 5000 parasites in 3 out of the 4 surviving colonies, was found in the untreated group in October, whereas a significantly lower mite infestation was recorded in the colonies where an appropriate acaricide treatment was carried out (Mann-Whitney U test: n_1_=4, n_2_=5, U=0, *P* = 0.007; Fig. 1a).

**Figure 1.**
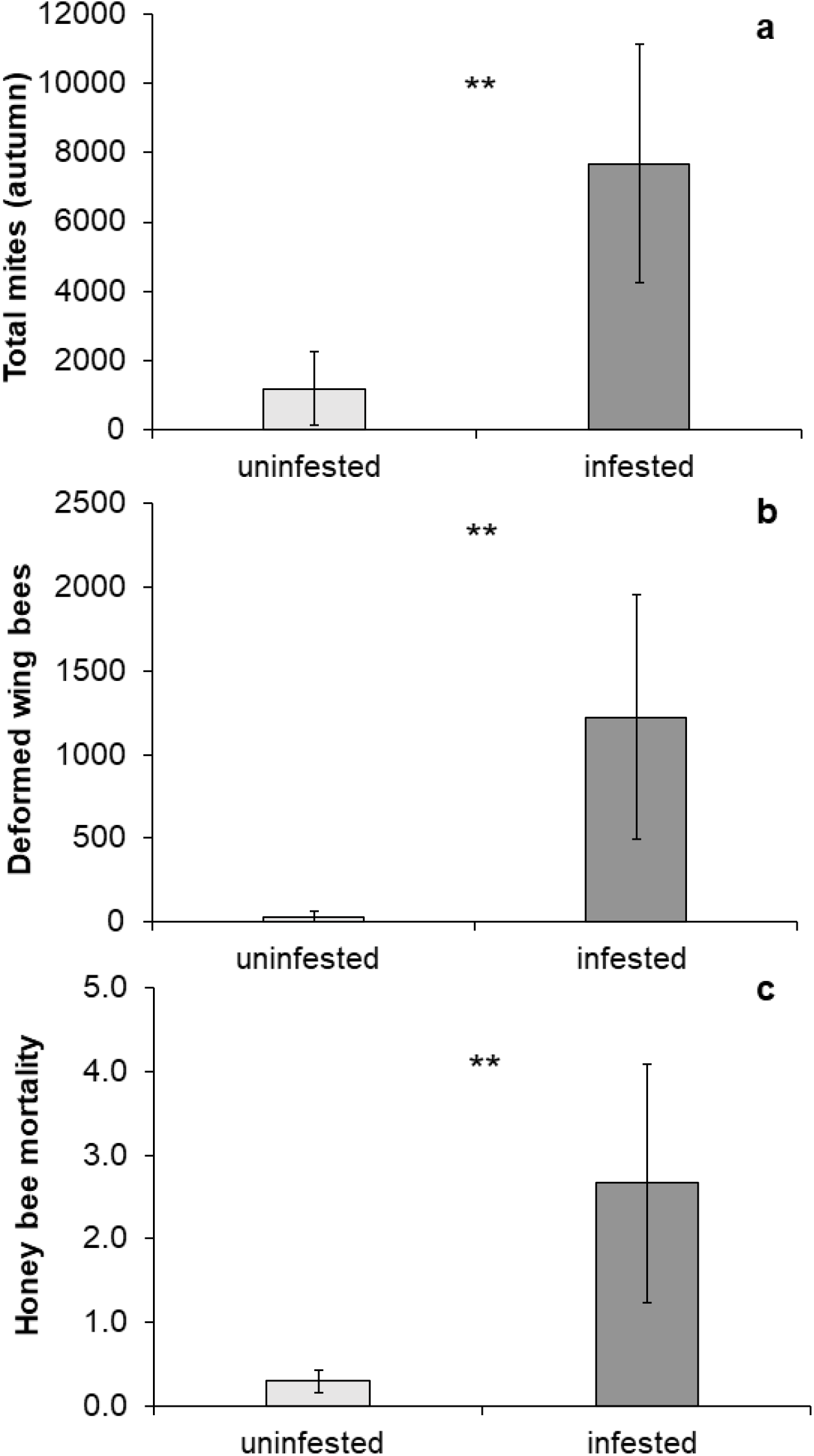
Effects of mite infestation on honeybee colonies: total mites collected from mite infested colonies and uninfested colonies after a control treatment (a); total number of deformed wing bees observed in mite infested and uninfested colonies along the experimental period (b); honey bee mortality (dead bees*1000/total bees*day) recorded in November in mite infested and uninfested colonies (c).

In turn, higher mite infestation caused increasing viral load in bees belonging to the colonies where the parasite population was higher. In fact, the proportion of individuals with deformed wings, a symptom associated to elevated viral infections, was higher in the colonies with higher parasitic pressure (Mann-Whitney U test: n_1_=5, n_2_=5, U=0, *P* = 0.005; Fig. 1b).

As a result of the increasing mite infestation and the associated viral infection, a higher bee mortality was observed in the group of colonies suffering from a higher parasitic pressure (Mann-Whitney U test: n_1_=4, n_2_=5, U=0, *P* = 0.007; Fig. 1c).

This in turn accelerated seasonal depopulation in mite infested colonies, such that in November a significantly lower number of bees was found in mite infested colonies as compared to uninfested ones (Mann-Whitney U test: n_1_=5, n_2_=5, U=0, *P* = 0.005; Fig. 2a). By October one mite infested colony had collapsed, soon followed by another two, whereas no losses were observed in the other group.

**Figure 2.**
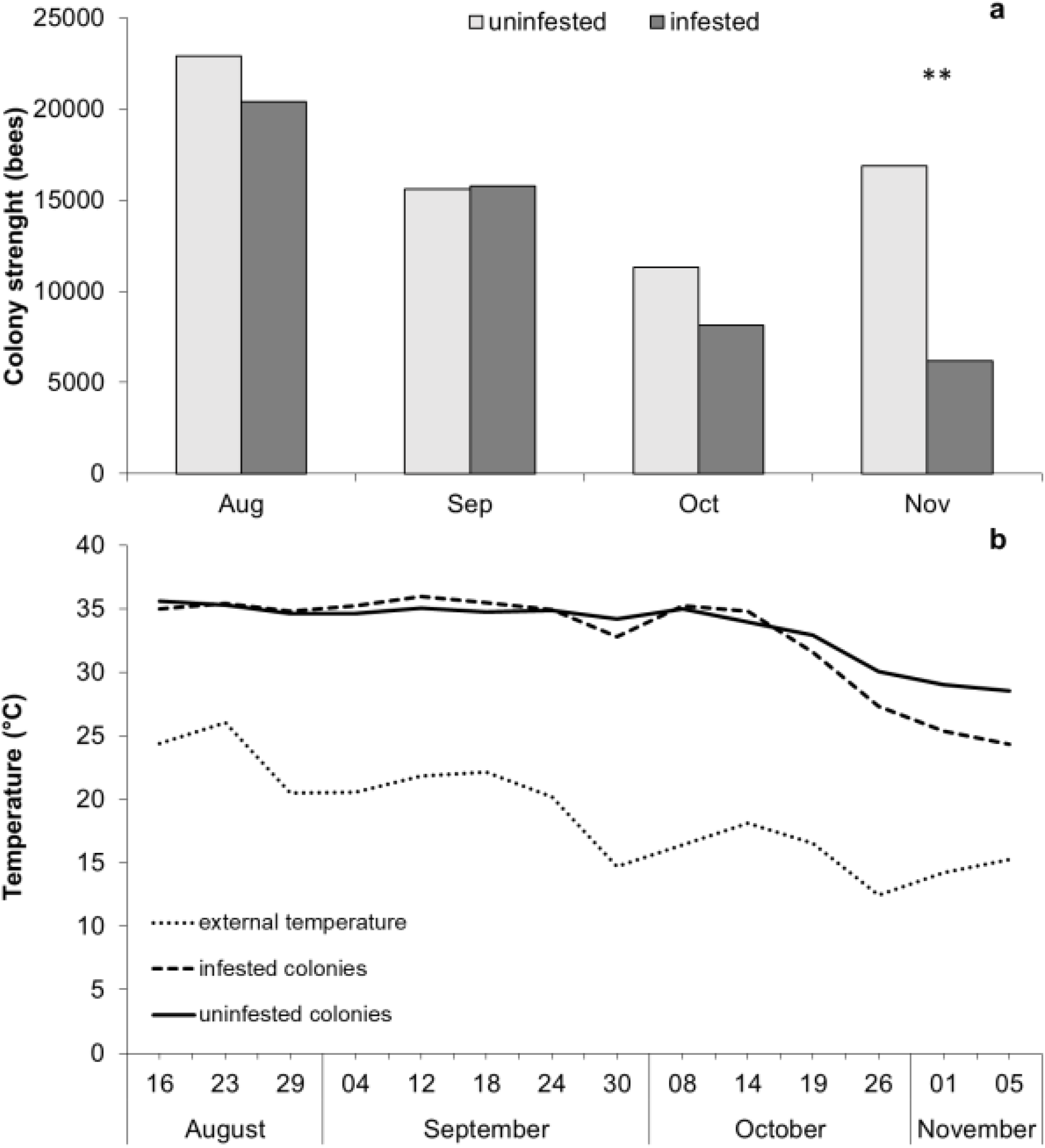
Estimated number of honey bees recorded during the trial in mite infested and uninfested colonies (a) and temperature recorded inside mite infested and uninfested colonies during the trial as compared to the external temperature (b).

### 2.2. Temperature control in mite infested colonies

The average daily environmental temperature gradually decreased from August, when 26 °C was recorded, to November, when it reached 15 °C (Fig. 2b). In the same period, the temperature inside the hives showed a concurrent, albeit less marked decrease, starting from 35.5 °C registered in August. However, whilst the internal temperature of control colonies was around 30 °C in November, that of mite infested colonies dropped to 25 °C in the same period (Fig. 2b).

### 2.3. The combined effect of low temperatures and mite infestation on individual bees

Since the field trial revealed that the temperature within the nest can be lower than optimal in both uninfested and mite infested colonies by a few Celsius degrees, we investigated how this can affect the survival of uninfested bees and bees that were mite infested during the pupal stage. To this aim, at eclosion, we exposed both mite infested and uninfested adult bees to different temperature regimes under laboratory conditions and assessed the effect on DWV replication, survival of bees and expression of some selected genes.

As expected, mite infestation significantly influenced viral replication, such that bees parasitized by one mite during development had a higher viral load as compared to unparasitized bees (two-way ANOVA test: d.f. = 1, F = 16.873, *P* < 0.001; Fig. 3) while low temperature did not affect viral dynamics (two-way ANOVA test: d.f. = 1, F = 2.799, *P* = 0.106; Fig. 3).

**Figure 3.**
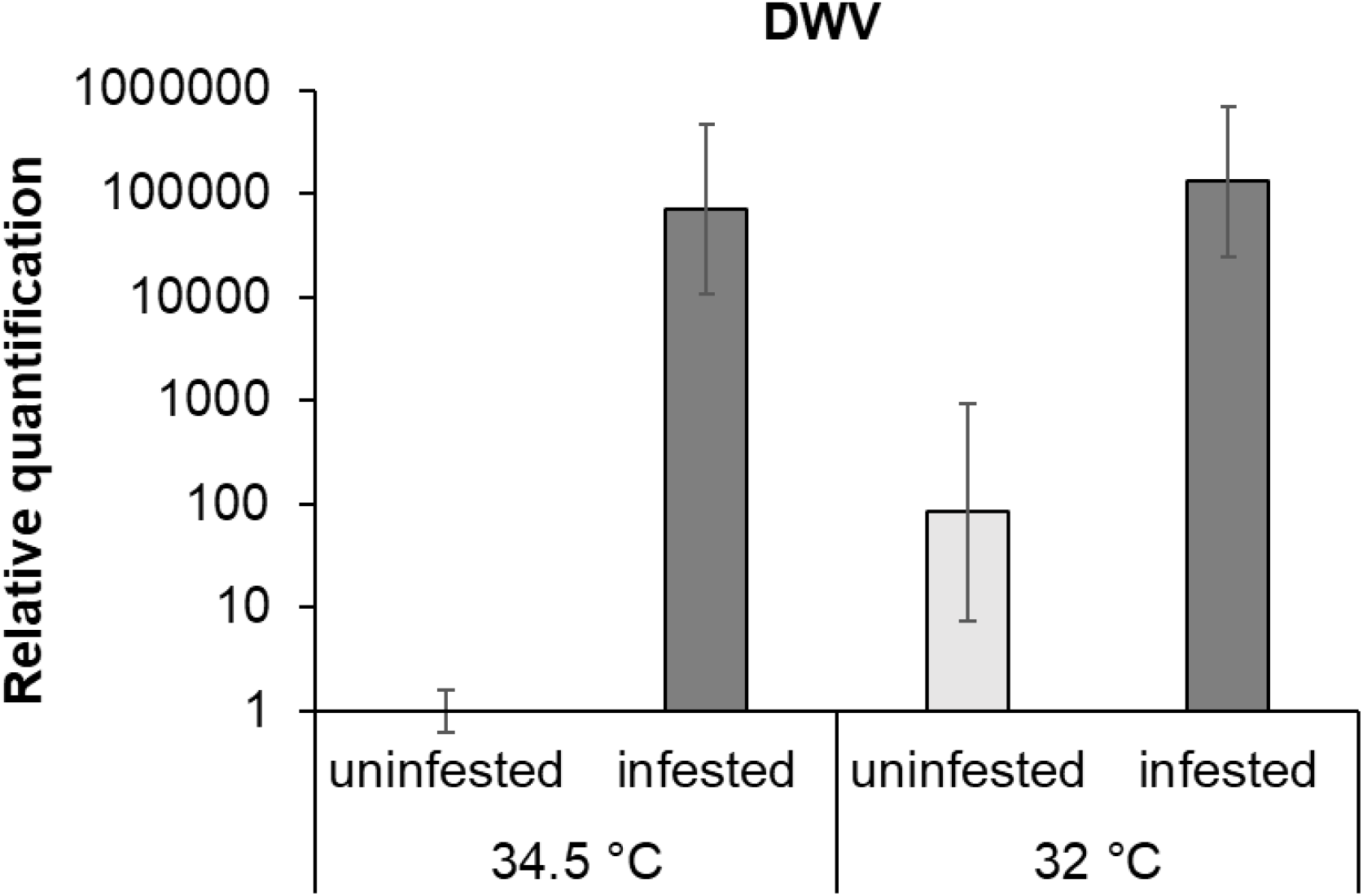
DWV infection in uninfested and mite infested bees exposed to two different temperatures. *Varroa* infestation is associated to higher virus load while low temperature and the interaction of mite and cold stress did not affect DWV load.

As a result of mite infestation and viral infection, a reduced survival of parasitized bees was observed (Cox regression analysis: HR=1.759, *P* < 0.001; Fig. 4); also, a similar but smaller effect of a lower rearing temperature was observed (Cox regression analysis: HR=1.274, *P* = 0.027; Fig. 4). Interestingly, mite infested bees exposed to low temperatures (i.e. 32 °C) had a longevity further reduced in comparison to control bees and bees exposed to either mite infestation or low temperature. Since the interaction between the infestation and low temperature is not significant (Cox regression analysis: HR=1.153, *P* = 0.516; Fig. 4) the observed reduced survival seems to be due to an addictive effect of the two stressors.

**Fig. 4.**
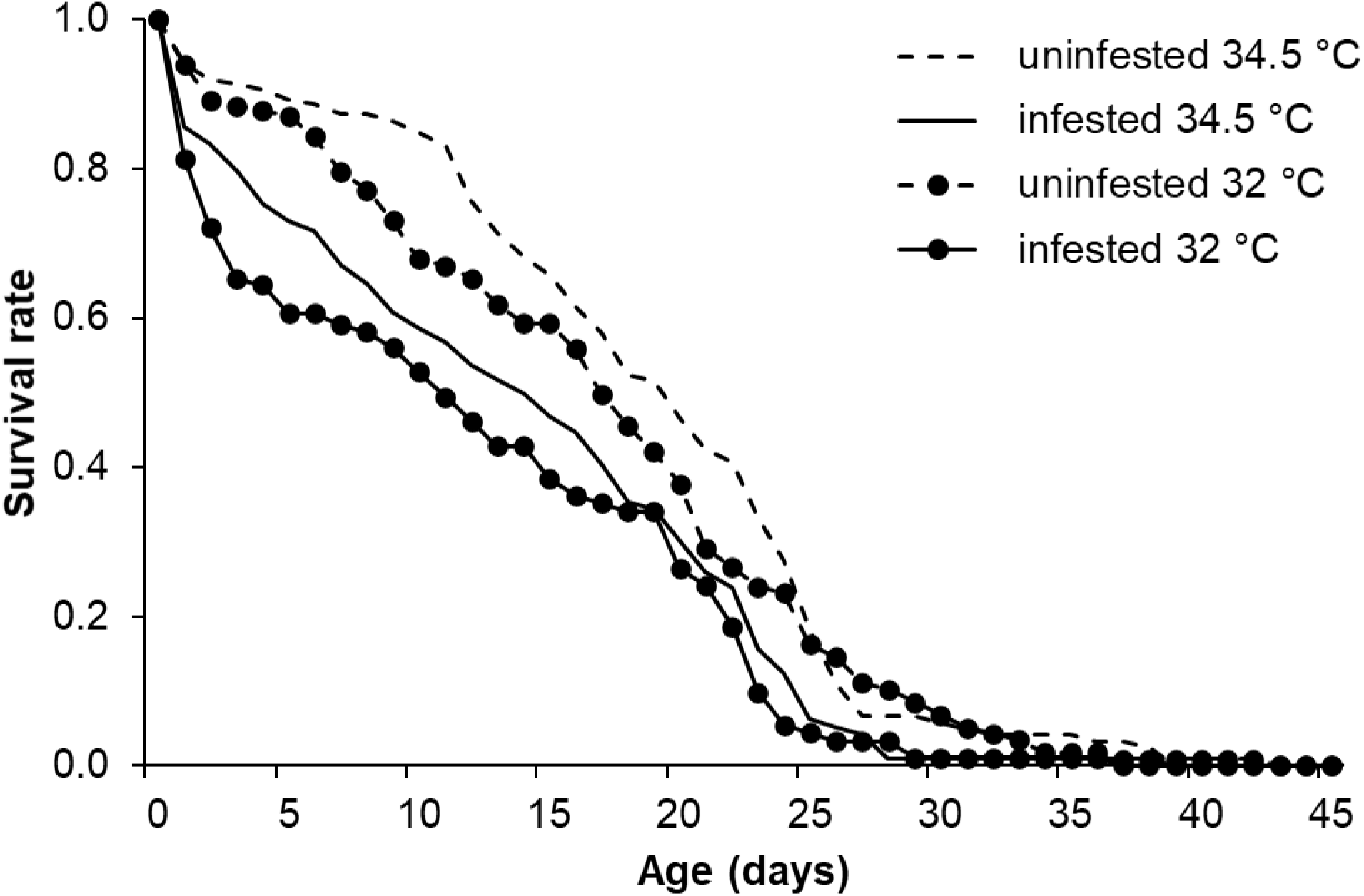
Survival of honey bees exposed to *V. destructor* during the pupal stage and exposed to two temperature regimes at the adult stage.

We also exposed bee pupae to different temperature regimes under laboratory conditions and assessed the effect on development and survival. We found that the low temperature (i.e. 32 °C) did not affect the survival of bee pupae (the proportion of eclosing bees from pupae maintained at 32 and 34.5 °C was 89.1 and 90.3%, respectively); however, all pupae maintained at 32 °C took 2 days longer than normal to reach the adult stage.

Since honey bees are capable of producing heat in response to low external temperatures by contracting their thoracic flight muscles, provided that a convenient food supply is available, we wondered if mite infested bees are as efficient as uninfested bees in this activity when exposed to a sub-optimal temperature. We found that bees infested at the pupal stage responded less efficiently to a lower temperature than uninfested bees (Mann-Whitney U test: n1 = 12, n2 = 12, U = 19, *P* = 0.001; Fig. 5).

**Figure 5.**
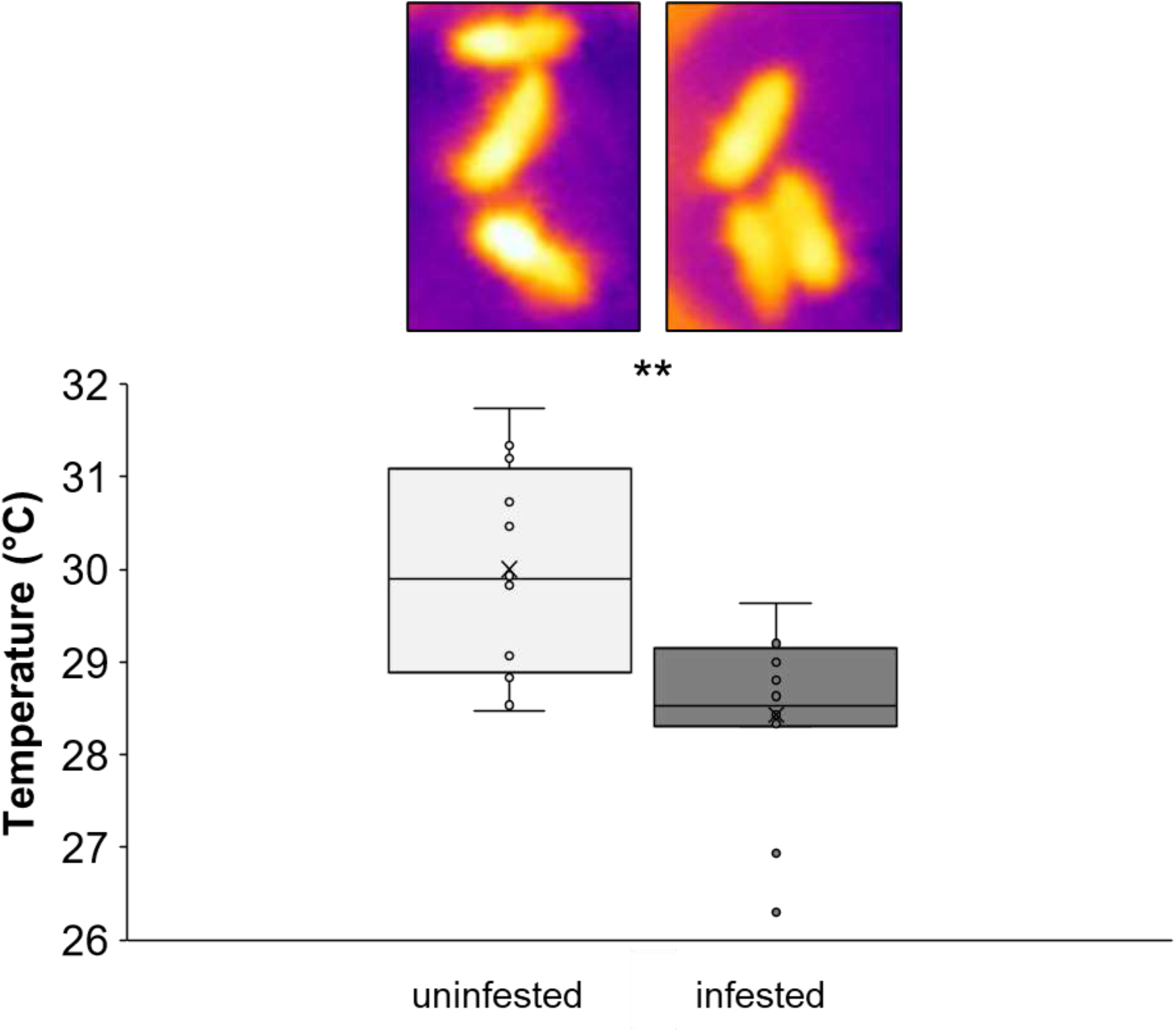
Average body temperature of mite infested and uninfested honey bees as assessed with a thermographic camera (FLIR, model i5).

The feeding activity of *V. destructor* during the pupal stage of bees, normally results in lighter emerging bees which often display underdeveloped wings, suggesting that some anatomical damage can occur because of the parasitic infestation. We therefore tested the effect of mite parasitization on the histology and mass of flight muscles in emerging bees. The flight muscles of bees exposed to mite infestation resulting in deformed wings did not appear to be affected in comparison to those of unparasitized bees (Fig. 6a). Furthermore, no significant differences were found between the weight of thorax (where flight muscles are located) of mite infested and uninfested bees (Mann-Whitney U test: n1 = 10, n2 = 10, U = 34, *P* = 0.113; Fig. 6b; Tab. S1). Instead, the different weight of parasitized bees appeared to be related to the lower amount of haemolymph that can be found in bees as a result of mite feeding (Fig. 6b, Tab. S1).

**Fig. 6.**
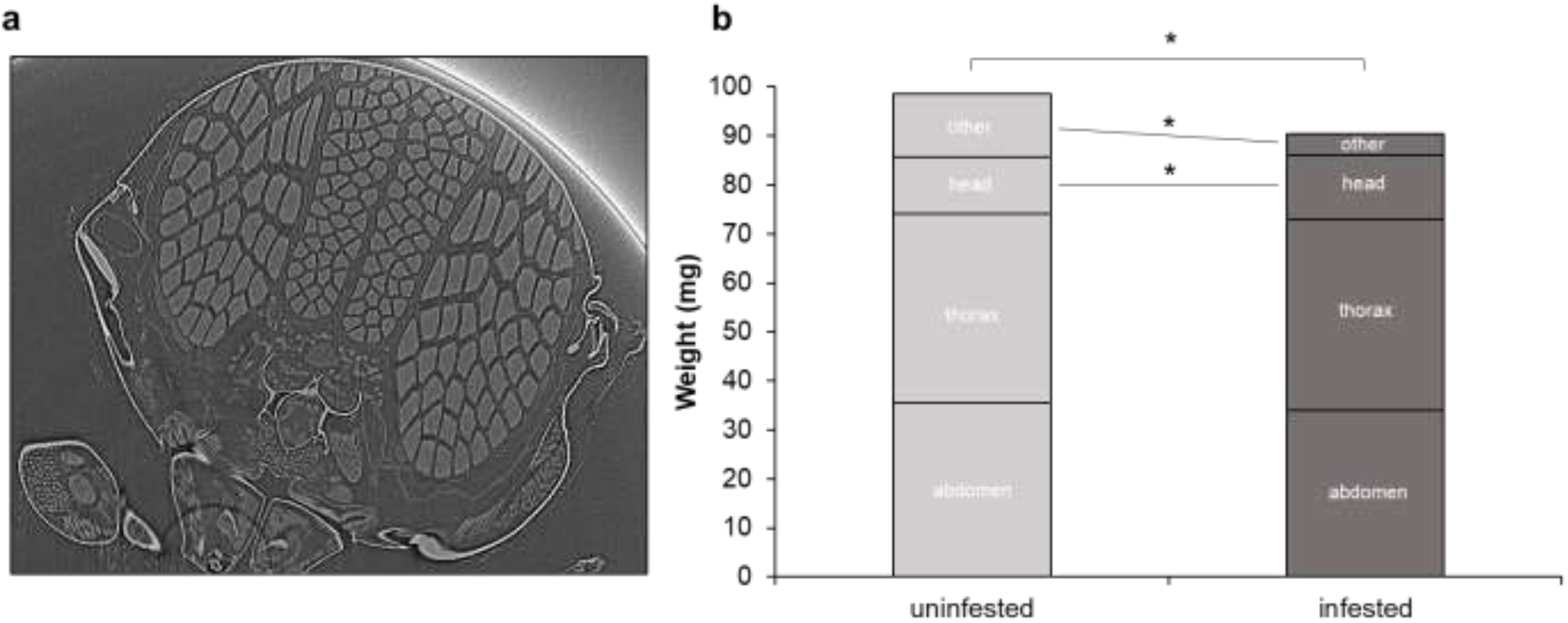
MicroCT scan of DWV symptomatic honey bee thorax (a). Weight of mite infested and uninfested eclosing honey bees and their body parts; “ other” likely represent the weight of the haemolymph lost after dissection (b).

After proving that no anatomical deficit was apparent in mite infested bees, we investigated if a sufficient amount of nutrients was available to support the metabolic activity of such apparatus and investigated sugar consumption in mite infested bees as compared to uninfested ones. As expected, we found that, at low temperatures, both uninfested and mite infested bees increased sugar consumption (two-way ANOVA test: d.f. = 1, F =7.412, *P* = 0.009; Fig. 7); however, in case of mite infestation, sugar consumption was significantly reduced (two-way ANOVA test: d.f. = 1, F = 21.09, *P* < 0.001; Fig. 7).

**Fig. 7.**
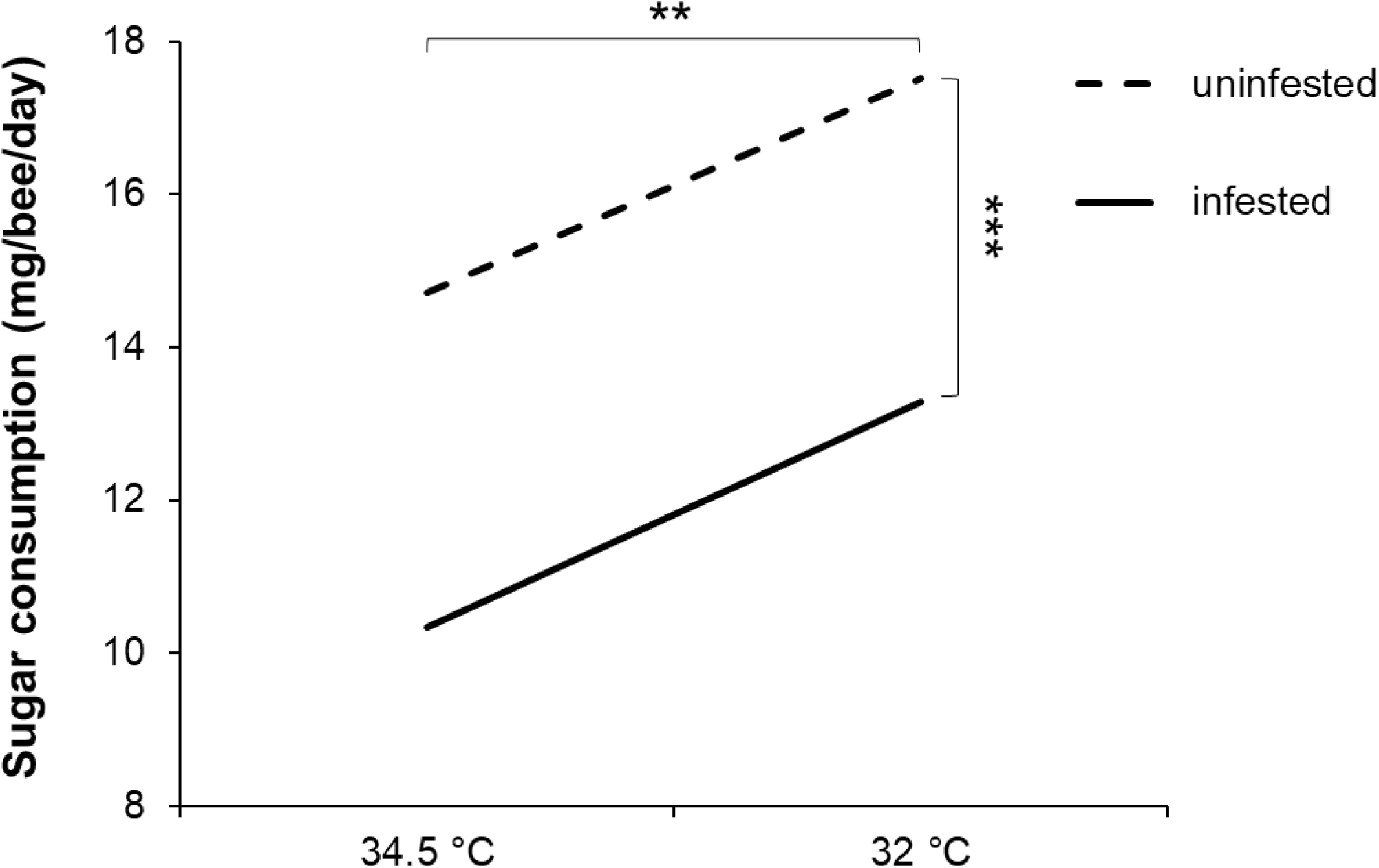
Sugar consumption of mite infested and uninfested bees exposed to two temperature regimes.

### 2.4 The effect of Varroa parasitization on insulin/insulin-like signalling (IIS) pathway

To gain insight into the *Varroa* induced anorexia on honey bees, we studied the expression of insulin receptor substrate 1 (IRS-1), a key protein in the insulin/insulin-like signalling (IIS) pathway as affected by mite infestation. IRS-1 appeared to be up-regulated in mite infested bees (Mann-Whitney U test: n1=12, n2=12, U=38, P = 0.025; Fig. 8a) highlighting an influence of *Varroa* mite infestation on bee metabolism. Furthermore, to verify that the upregulation of IRS-1 in mite infested bees was not influenced by the reduced dietary input in those bees, we studied IRS-1 expression in uninfested bees reared at the standard temperature of 34.5 °C and at 32°C, a condition under which an increased sugar consumption is observed. In this case, relative expression of IRS-1 was not different between bees maintained at normal and lower temperature (Mann-Whitney U test: n1=8, n2=10, U=34, P = 0.297; Fig. 8b).

**Fig. 8.**
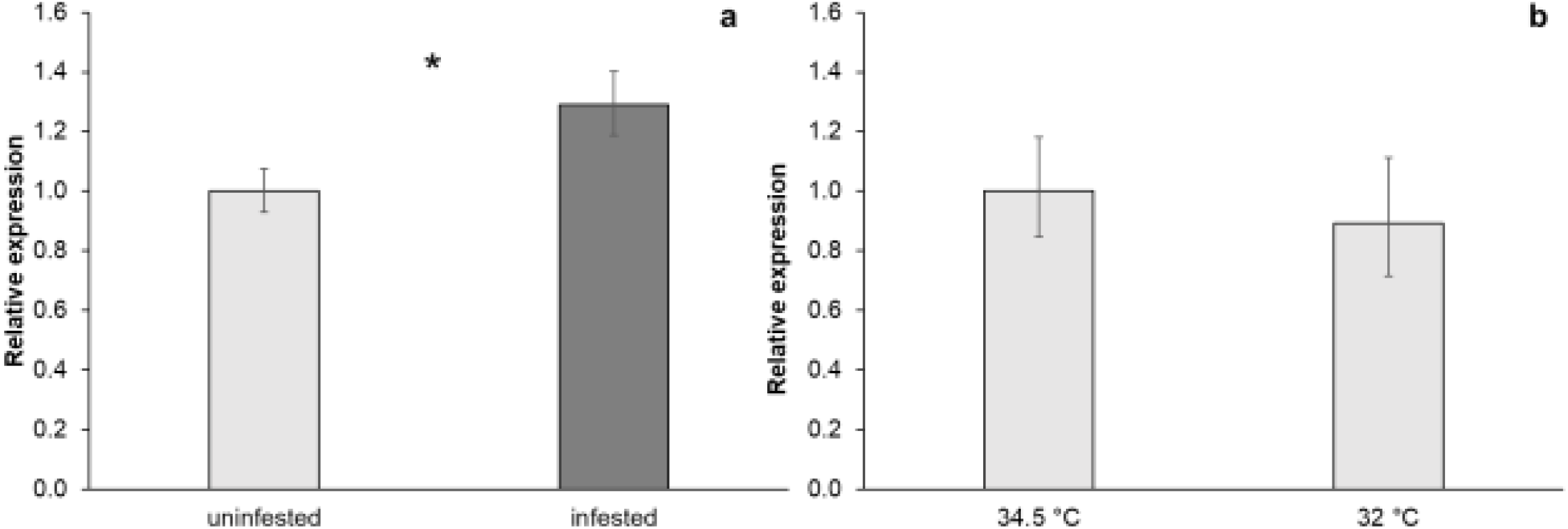
Relative expression of IRS-1 in bees infested or not with *Varroa destructor* at 34.5 °C (a) and in bees reared at 34.5 °C and 32 °C (b).

## 3. Discussion

The progressive decline of mite infested honey bee colonies towards the end of Summer is a very common situation under temperate climates in the Northern Hemisphere and was clearly confirmed here. In particular, we provided further evidence that this decline is related to the increased mortality of bees associated to high DWV infection levels caused by the parasitic activity of the mite *V. destructor*, vectoring the virus and triggering its replication in infected bees^17,26,27^.

We also showed that colony decline parallels the concurrent decrease in environmental temperature, which is observed in the Autumn months under these climatic conditions. Moreover, we noted that mite infested honey bee colonies are less efficient at countervailing the decreasing temperature, as sub-optimal temperatures are more often observed in those colonies during this period. This is likely due both to the reduced number of bees in mite infested colonies and to the reduced capacity of the surviving bees to thermoregulate, as demonstrated under laboratory conditions.

A detailed laboratory investigation into the effects of the concurrent exposition of bees to a parasitic infection and sub-optimal temperatures, clearly showed that the negative effects of these two stressors add up to reduce the survival of bees. In particular, by recording the temperature of both mite infested and uninfested adult bees upon exposition to a low temperature, we noted that the first have a reduced capacity to warm up their bodies to counteract the lowering external temperature: a result that, to our knowledge, has not been reported before, despite the thermoregulatory capacity of bees has been investigated in considerable detail^28,29^.

Our results indicate that the flight muscles of parasitized bees are normally developed. A great deal of research has been dedicated to the effect of mite infestation on individual bees; however, so far^30,31^, apart from some early studies on wing development^32^ and some recent results regarding the fat body beneath the feeding hole^33^, little is known about the anatomy of honey bees affected by mite parasitization. Here we suggest that the internal anatomy of infested bees’ thorax is not impaired by the feeding mite.

On the other hand, we noticed that the sugar intake of mite infested bees is significantly affected, suggesting that mite infested bees suffer from a kind of parasite induced anorexia. In addition, the expression pattern of IRS-1, a key gene in the IIS pathway, suggests that this anorexia is probably elicited by the molecular effects of *Varroa* parasitization on bee’s metabolism. In fact, this protein is upregulated in presence of the mite but is not modified by sugar intake; this latter evidence confirming previous results about the lack of response of insulin receptor substrate and insulin like peptide 2 to dietary inputs^34^. Recent results showing that the downregulation of insulin receptor 1, stimulates the sensitivity to sucrose and weight gaining by honey bees^35^ seem to confirm our proposed mechanistic explanation of the observed anorexia.

In the honey bee the insulin/insulin-like signalling pathway is involved in lifespan and regulation of social behaviors, including the onset of foraging and the type of collected food^36,37,38^. The regulation of behavioral maturation has a strong influence on the total lifespan of honey bees with those commencing foraging at a younger age living significantly shorter than those foraging later^39^. Since IIS genes are differentially expressed in forager and nurse bees^37^, it is possible that the previously reported accelerated behavioral maturation in mite infested bees^40,41^ could be responsible for the effect of *Varroa* on IIS pathway. In any case, the mite induced anorexia we report here parallels other cases of disease induced anorexia in insects, and in particular, the reduced feeding of caterpillar larvae infected by nucleopolyhedrovirus (NPV)^42^. Furthermore, we observed that the lower temperature inside the hive could also affect the developmental time and the survival of emerging adult bees.

In conclusion, it appears that mite infestation further than increasing *per se* the mortality of bees, reduces the capacity of bees to thermoregulate, exposing them to the detrimental effect of lower temperatures. In addition, the reduction in the number of bees engaged in thermoregulation together with their reduced efficiency, can in turn affect the developing bees further aggravating the phenomenon. In sum, a number of dangerous positive feed-back loops are generated, with devastating effects on the survival of the colony, clearly revealed by our field results. Therefore, it appears that the decreasing temperature observed during the cold season can enhance the negative effect of the increasing mite infestation, further reducing the survival of bees and thus impairing the very sustainability of the colony.

These results eventually provide a more comprehensive understanding of the reported higher colony losses in northern regions^12,13,14^ where lower temperatures are observed during the cold season^16^. Furthermore, these results may provide additional insight into the peculiar world distribution of colony losses. In 2010, Neumann and Carreck^2^ noted that, until that date, large losses of honey bee colonies had been observed only north of a parallel separating the regions of the world where the mite *V. destructor* had been responsible of severe infestations from those where the mite, albeit present, had had a reduced impact. They therefore coined the fortunate expression “ *Varroa* equator” to denote that parallel, underlining the crucial role of the mite in the phenomenon of colony losses. Further studies clearly indicated that the ultimate responsible of the loss of bee colonies observed in the northern hemisphere is the deformed wing virus, which has a worldwide distribution^43,44^. Here we would like to further refine the concept of “ *Varroa* equator” suggesting that the occurrence of elevated colony losses north of that parallel is certainly the result of the impact of the *Varroa*-DWV association on bee colonies but this impact is further exacerbated by the lower temperatures that are observed in the northern hemisphere in the Autumn months; a situation that is certainly rarer in the Southern hemisphere where most land masses are distributed north of the 40^th^ parallel where milder winters are more common. Finally and importantly, the results reported here highlight the relevant role that abiotic factors can have in shaping the effect that biotic factors already exert on the health of honey bees. To our knowledge this is the first study, in honey bees and insects in general, dealing with the interaction between an abiotic factor such as the environmental temperature and a widespread parasite. Although some conclusions of this study are clearly restricted to honey bees with their peculiar biology, the experimental approach adopted here (i.e. multilevel and holistic) can represent a useful template for similar studies on other insect species, aiming at elucidating the critical positive feed-back loops triggered under these conditions. We hope that similar studies will become more common in view of the alarming news regarding climate change and its potential impact on terrestrial ecosystems^45^ and the continuous pressure of natural and exotic parasites on insect populations^46,47^.

## 4. Materials and methods

### 4.1. Mite infestation and temperature control in the hive

Two apiaries, made of 5 colonies each, housed into ten frames Dadant Blatt hives (385 mm x 452 mm x 310 mm), were set up in Udine (Italy); in particular, the untreated apiary was placed north of the city (46°04’54.2” N, 13°12’34.2” E), while the treated one in south of it (46°02’10.9” N 13°13’24.9” E), about 4 km apart. Previous studies indicated that local colonies are hybrids between *Apis mellifera ligustica* Spinola and *Apis mellifera carnica* Pollmann^48,49^.

In the untreated apiary, no acaricidal treatments were carried out throughout the Summer, so that mite infestation could naturally increase along the season. In the treated apiary, mite population was kept under control with different acaricides; in particular, 2 strips of an Amitraz based product (Apitraz, Laboratorios Calier S.A.) were used in August, followed by three treatments with a thymol based product in tablets (ApiLife Var, Chemical Laif S.p.A) from September to mid-October and with 2 strips of a Fluvalinate based product (Apistan, Vita Europe) in October. Bee population in the experimental hives was estimated, approximately once a month from August to December, by counting the number of full or partial “ sixth of frames” covered by bees in each hive at sunset, considering that a fully covered sixth of comb corresponds to 253 adult bees^50^. The number of brood cells was estimated by means of the same method, considering that one sixth of frame of brood cells corresponds to 728 worker brood cells.

During the experiment, infestation levels were estimated by counting the number of mites fallen on a vaseline coated bottom board. At the end of the first decade of October, highly-infested hives of the untreated apiary were treated with 2 strips each of a fluvalinate based product. The total number of mites fallen in the next fifteen days confirmed the higher infestation level of the untreated hives (Fig. 1a). To assess bee mortality, dead bees found in under basket cages placed in front of the colonies were counted every week from August to November, taking note of the number of the individuals showing deformed wings.

Data collected from the field (i.e. infestation level, proportion of deformed bees, bees’ mortality and decrease of bee population) were analysed using a Mann-Whitney U test.

### 4.2. Temperature measurement in apiary

In order to monitor the temperature inside the hives, a temperature probe (Maxim integrated, US) was inserted in the central part of each hive during the warmer months (August -September) and then moved following the honey bee cluster (October - November); monitoring took place from August to November 2018. At the end of the experiment, data collected by the probes were transcribed and the average daily temperature with variability coefficient (standard deviation/mean) calculated.

Average daily external temperature data were derived from the regional meteorological observatory (ARPA FVG – OSMER and GRN, http://www.meteo.fvg.it/).

### 4.3. Artificial rearing of bee larvae under different temperatures

Two brood frames containing cells sealed by workers in the preceding 15 hours were collected from one hive of the experimental apiary of the Dipartimento di Scienze AgroAlimentari, Ambientali e Animali, University of Udine. Brood frames were transferred into the lab, uncapped and maintained in a climatic chamber (34.5 °C, 75% R.H.), until the 5^th^ instar larvae emerged from the brood cells; larvae were collected and transferred into 10 cm large Petri dishes. Half of the larvae were stored in a climatic chamber at 34.5 °C, 75% R.H. and half in a second chamber at 32 °C, 75 % R.H. Developing larvae were daily monitored and death individuals removed. The experiment was replicated three times. In total 379 larvae were used for this experiment.

### 4.4. Artificial infestation of bees

Honey bees and mites were collected from the same experimental apiary cited above. The mites and last instar bee larvae were collected from brood cells capped in the preceding 15 h obtained as follows. In the evening of the day preceding the experiment the capped brood cells of several combs were marked. The following morning the combs were transferred to the lab and unmarked cells, that had been capped overnight, were unsealed. The comb was then placed in an incubator at 34.5 °C and 75% R.H., where larvae and mites spontaneously emerged. Last instar bee larvae were transferred into gelatin capsules (Agar Scientific ltd., 6.5 mm diameter) with no mites (uninfested bees) or 1 mite (infested bees) and maintained at 34.5 °C, 75% R.H. for 12 days^51^.

### 4.5. Honey bee rearing under artificial conditions

Upon eclosion, newly emerged adult bees were separated from the infesting mite and transferred into four plastic cages (185 × 105 × 85 mm) with water and sugar candy (Apifonda®), supplied *ad libitum* through a small plastic container (Ø = 1.5 cm), refilled every 2 days and placed on the floor of the cages. To prevent the exsiccation of the candy, containers were wrapped with laboratory film (Parafilm®); a small cut was made on the top, to ensure bee feeding. Two cages, one with bees that were mite infested at the pupal stage and the other with the same number of uninfested bees were maintained in a climatic chamber at 34.5 °C, 75% R.H.; the other two cages were maintained in a climatic chamber at 32 °C, 75% R.H.; each cage hosted from 20 to 25 adult bees. Bee survival and diet consumption were recorded daily.

Seven days after the eclosion, at least eight bees per treatment were collected, killed with liquid nitrogen and stored at -80 °C until analysis. The experiment was repeated six times from July to September. In total, from 112 to 134 bees per group were used.

The hazard ratio was calculated by means of a weighted cox regression hazard model^52^ using R, version 3.6.2^53^.

Sugar candy intake under different conditions was monitored from day 3 after the emergence of bees to day 15 and normalized according to the different weights of infested and control bees^54^. Sugar consumption under different treatments was normalized and analysed by means of a two-way ANOVA test.

### 4.6. Honey bee thermoregulation

To test if mite infested bees thermoregulate as well as uninfested bees, honey bee larvae were artificially infested using the protocol described above^51^ or maintained uninfested as a control. Upon eclosion, newly emerged adult bees were separated from the infesting mite and transferred into plastic cages (185 × 105 × 85 mm). Uninfested and mite-infested honey bees were maintained in a climatic chamber at 34.5 °C, 75% R.H.

Starting from day 4, three honey bees collected randomly from the two groups (mite-infested and uninfested) were placed in polystyrene box, transferred to room temperature (25 °C) and photographed with an infrared thermographic camera (brand: FLIR; model: i5; thermal resolution = ± 0.1 °C) with emissivity settled at 0.97^55^. Pictures were taken through a hole in the polystyrene lid to reduce the possible interference of light radiation. Pictures were taken for four consecutive days with three technical replicates (i.e. three pictures) for each time point. Images were analysed with FLIR Tools® software and temperature data were collected, considering the average value of the warmer part of the bee which always corresponded to thorax. The area used to calculate the mean temperature was equal for each bee.

The recorded temperatures were compared using the Mann-Whitney U test.

### 4.7. Honey bee weight

To check the condition of flight muscles of experimental bees, we artificially infested or not honey bee larvae as described before. The two groups of bees (uninfested and infested) were maintained in a climatic chamber at 34.5 °C, 75% R.H., dark, for 12 days. Upon eclosion, newly emerged honey bees were weighted and dissected into head, thorax and abdomen; then, each section was separately weighted.

The weight of the different body parts of mite infested and uninfested bees was compared with the Mann-Whitney U test. In total, 10 uninfested and 10 mite infested bees were used.

### 4.8. Gene expression analysis and DWV quantification

Sampled bees were defrosted in RNA*later* (Ambion®) and gut deprived. The whole body of the bees was homogenized by means of mortar and pestle in liquid nitrogen. Total RNA was extracted from each bee, according to the method suggested by the producer of RNeasy Plus mini kit (Qiagen®, Germany). The amount of RNA in each sample was quantified using a NanoDrop® spectrophotomer (ThermoFisher™, US) and integrity verified by means of agarose gel electrophoresis. cDNA was synthetized starting from 500 ng of RNA following the manufacturer specifications (PROMEGA, Italy). Additional negative control samples containing no RT enzyme were included. Ten ng of cDNA from each sample were analysed by qRT-PCR with the primers reported in Table 1, using SYBR®green dye (Ambion®), according to the manufacturer specifications, on a BioRad CFX96 Touch™ Real time PCR Detector. Primers efficiency was calculated according to the formula E=10^(−1/slope)-1)*100. The following thermal cycling profiles were adopted: one cycle at 95 °C for 10 minutes, 40 cycles at 95 °C for 15s and 60 °C for 1 min, and one cycle at 68 °C for 7 min.

**Table 1.**
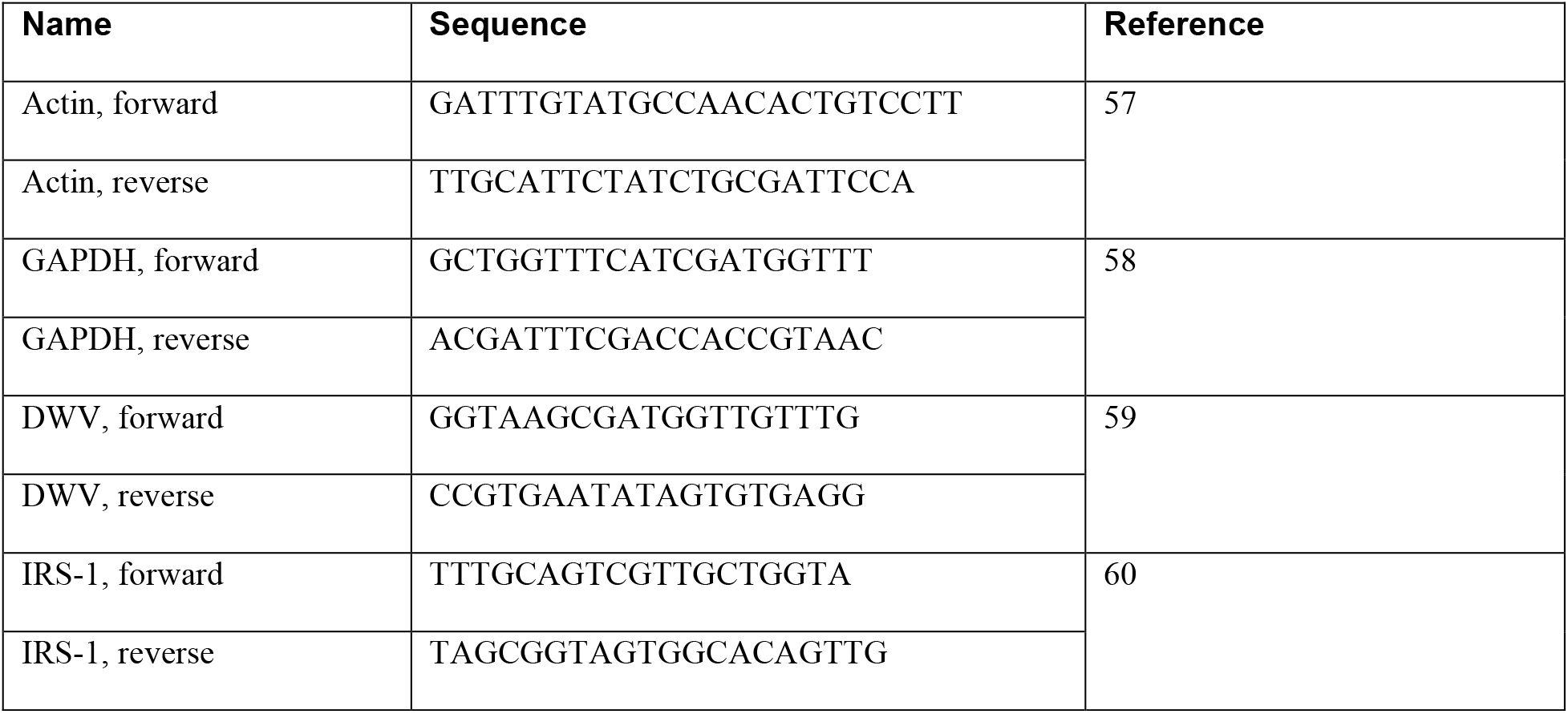
Primer pairs used in this study.

Relative gene expression data were analysed using the 2-ΔΔCt method^56^ using actin and GAPDH as housekeeping genes. Relative expression data were transformed, normalized and analysed by means of two-way ANOVA test performed with R statistical software, version 3.6.2. (R Core Team, 2019). At least eight individual bees per experimental group were analysed.

### 4.9 IRS-1 expression in mite infested and uninfested honey bees

To study IRS-1 expression level in mite infested bees, we artificially infested or not honey bee larvae as described before. The two groups of bees (uninfested and mite infested) were maintained in a climatic chamber at 34.5 °C, 75% R.H., dark, for 12 days. Upon eclosion, mite infested and uninfested newly emerged bees were reared separately in plastic cages. At day 7 bees were sampled to assess the expression of the selected gene. The protocols used for RNA extraction, cDNA synthesis and qRT-PCR are the same as those described above. Relative expression data were analysed by means of the Mann-Whitney U test. Three replicates of the experiment were performed. Four bees per replicate were analysed.

## 5. Acknowledgements

This research was funded by the European Union’s Horizon 2020 research and innovation programme, under grant agreement No 773921 (PoshBee) and by the Italian Ministry of University, PRIN 2017 - UNICO (2017954WNT). The microCT scan images were carried out at Elettra Synchrotron Light Laboratory in Trieste (Italy) within the framework of proposal n. 20190281. Authors would like to thank Prof. Alessandro Peressotti for the concession of the thermographic camera used in this article.

## 6. Author contributions

D.A. D.F. F.N. designed the research; L.A., D.A., G.B., M.D.A., S.D.F., D.F., F.N., V.Z. performed the research; D.A., D.F., F.N., V.Z., analyzed the data; D.A. D.F. F.N. wrote the paper. All authors revised the final version of the manuscript.

## Notes

### Competing Interest Statement

The authors have declared no competing interest.

### Summary of Updates

To gain insight into the Varroa induced anorexia on honey bees, we further studied the expression of insulin receptor substrate 1 (IRS-1), a key protein in the insulin/insulin-like signalling (IIS) pathway as affected by mite infestation.

